# Conserved sites on the influenza H1 and H3 hemagglutinin recognized by human antibodies

**DOI:** 10.1101/2024.10.22.619298

**Authors:** Daniel P. Maurer, Mya Vu, Ana Sofia Ferreira Ramos, Haley L. Dugan, Paul Khalife, James C. Geoghegan, Laura M. Walker, Goran Bajic, Aaron G. Schmidt

**Author notes:** **Correspondence:** Aaron G. Schmidt. These authors contributed equally (Mya Vu, Ana Sofia Ferreira Ramos, Haley L. Dugan, Paul Khalife).

## Abstract

Monoclonal antibodies (mAbs) targeting the influenza hemagglutinin (HA) have the potential to be used as prophylactics or templates for next-generation vaccines that provide broad protection. Here, we isolated broad, subtype-neutralizing mAbs from human B cells targeting the H1 or H3 HA head as well as a unique mAb targeting the stem. The H1 mAbs target the previously defined lateral patch epitope on H1 HAs and recognize HAs from 1933 to 2021 in addition to a swine H1N1 virus with pandemic potential. Using directed evolution, we improved the neutralization potency of these H1 mAbs towards a contemporary H1 strain. Using deep mutational scanning of four antigenically distinct H1N1 viruses, we identified potential viral escape pathways. For the H3 mAbs we used cryo-EM to define the targeted epitopes: one mAb recognizes the side of the H3 head, accommodating the N133 glycan and a pocket underneath the receptor binding site. The other H3 mAb recognizes an epitope in the HA stem that overlaps with previously characterized mAbs, but with distinct antibody variable genes and mode of recognition. Collectively, these mAbs identify common sites recognized by broad, subtype-specific mAbs that may be elicited by next-generation vaccines.

## INTRODUCTION

Two influenza types cause human disease: influenza A viruses (IAVs) emerge from animal reservoirs and cause global pandemics while influenza B viruses (IBVs) only circulate in the human population. Antigenic and genetic differences of the major surface glycoprotein hemagglutinin (HA), which engages the sialic acid receptor and fuses the viral and endosomal membranes, further divide IAVs into two groups with 18 HA subtypes (H1–H18) total. In the last century, IAVs bearing H1 or H2 (group 1), or H3 (group 2) HAs have circulated in the human population; H1N1 and H3N2 viruses currently circulate.

Vaccine effectiveness against seasonal influenza viruses reaches modest levels around 50%^1^ and updated vaccines are needed due to antigenic drift – the acquisition of amino acid mutations in HA that preserve function but escape population-wide antibody recognition. IAVs undergo antigenic drift more rapidly than IBVs^2^, requiring continuous monitoring for new antigenic variants. Additional barriers to an effective vaccine include complex and variable immune histories within the human population. A complementary approach to seasonal vaccination is to identify monoclonal antibodies (mAbs) that recognize conserved HA sites for prophylactic use, particularly in high-risk populations. Characterizing such mAbs can also serve as templates to guide next-generation vaccine designs and inform how viral escape from vaccine-induced immunity may occur.

Antibodies that target the HA “head” and prevent sialic acid receptor engagement are an established correlate of protection but are usually subtype-specific^3–5^. mAbs with heterosubtypic breadth typically engage epitopes on the membrane proximal HA “stem” within group 1 or group 2, and, in some cases, both^6–10^. Although these stem-directed mAbs are protective in mice^6–10^, they require Fc-effector functions for protection, suggesting that although stem-directed mAbs can neutralize *in vitro*, there is minimal neutralization activity *in vivo*^11^. Due to their reactivity to potentially pandemic subtypes of IAVs that have not circulated in humans, stem-directed mAbs may help protect from severe disease in the case of zoonotic spillover or viral escape from anti-head mAbs. However, while passive transfer of stem-directed antibodies offers some protection, head-directed antibodies better protect against acquiring infection even when normalized by binding titers^12^.

In contrast to anti-stem mAbs, IAVs escape anti-head antibodies frequently at the population level, demonstrating that these mAbs provide a significant barrier for transmission that the virus must overcome to persist in the population^2,13,14^. Although epitopes in the head are more variable than the stem, it is nevertheless possible to identify broad, subtype-specific anti-head mAbs^15–24^. Epitopes on the side of the HA head (*e.g.,* the lateral patch, vestigial esterase) are conserved^15,21,22^, affording mAbs targeting these sites considerable breadth and potency. If next-generation vaccines can elicit antibodies targeting these conserved epitopes^25–27^, it is unclear how such a diverse polyclonal response would structurally recognize HA and ultimately how the virus would evolve to escape.

Here, we isolated and characterized broad H1 and H3 human mAbs capable of neutralizing human IAVs spanning several decades. For the broad H1 mAbs, which neutralize H1N1 IAVs spanning ∼100 years, we used four deep mutational scanning H1N1 viral libraries to map critical residues within the targeted epitope. Although escape from broad H1 mAbs was readily selected *in vitro*, human influenza H1N1 isolates have rarely contained these escape mutations. Broad H1 mAb neutralizing potency was improved against a contemporary H1N1 strain using directed evolution. For the broad H3 mAbs, which neutralize human H3N2 IAVs spanning ∼50 years, we determined cryo-EM structures in complex with HA. One broad H3 mAb targets an epitope in the head domain with regions of low mutational tolerance^28^. The other broad H3 mAb targets the group 2 stem site but engages with a distinct mode of recognition. The epitopes identified by these broad H3 mAbs overlap with those targeted by previously characterized mAbs, but with different antibody variable gene usage and paratopes. These data suggest that while antibodies targeting these conserved sites could produce viral escape variants, mAbs targeting these sites may nevertheless serve as prophylactics for high-risk populations.

## RESULTS

### Isolation of broad H1 and H3 monoclonal antibodies

To identify broad subtype neutralizing mAbs, we used a panel of reporter IAVs spanning 1933 to 2021 that includes distinct antigenic clusters^2^, representing IAVs that have previously escaped humoral immunity. We sorted single H1-reactive B cells from human PBMCs using recombinant A/California/07/2009 HA (**Supplementary Fig. 1A**). We then sequenced the variable genes, recombinantly produced mAbs^29^, and screened for neutralization activity. H1-targeting mAbs were screened against representative H1N1 strains A/WSN/1933, A/Solomon Islands/3/2006, and A/Michigan/45/2015. We identified two clonally related mAbs, ADI-77470 and ADI-77474, that completely neutralized (>99%) all three H1N1 viruses at 1 µg/ml used for screening (**Supplementary Fig. 1B**).

To isolate broad H3 mAbs, we sorted B cells using recombinant HA probes from H3N2s A/Hong Kong/1/1968 (HK68) and A/Darwin/6/2021 (DAR21) coupled to different streptavidin fluorophore conjugates to identify cross-reactive B cells (**Supplementary Fig. 1C**). The HK68 probe included the HA head appended to a foldon trimerization tag, allowing us to isolate head-directed mAbs in addition to those that target full-length DAR21 HA. The frequency of cross-reactive HK68/DAR21 head-reactive memory B cells was rare, ranging from 0.0049 to 0.016% across five donors (**Supplementary Fig. 1C**). We further characterized two H3 mAbs, ADI-85647 and ADI-85666 after screening for broad neutralization activity (**Supplementary Fig. 1D**).

We characterized the subtypic neutralization potency and breadth of these broad H1 and H3 mAbs against 11 H1N1 or 10 H3N2 IAVs. The broad H1 mAbs potently neutralized all H1N1 viruses, but not a representative H3N2; the broad H3 mAbs neutralized all H3N2 viruses, but not a representative H1N1 (**Fig. 1**), demonstrating breadth within but not across subtypes. To further test the breadth of these mAbs, we screened for binding by ELISA to the HA from a swine H1N1 with pandemic potential (A/Swine/HN/SN13/2018^30^) or an H3 HA from an avian-origin canine IAV found to evolve mammalian influenza-like traits (A/Canine/Hainan/79/2019^31^). Although these IAVs have never circulated in humans, these mAbs bound these strains, highlighting their considerable breadth (**Fig. 1C**). We performed competition assays to crudely map the region on HA targeted by these mAbs. The broad H1 mAbs competed with 047-09M-2F03^17^, a mAb that engages the lateral patch of H1 HAs. ADI-85647, a broad H3 mAb, competed with F005-126^32^, H3v-47^15^, and c585^21^, which target the H3 head, whereas the other H3 mAb, ADI-85666, competed with MEDI8852^9^, showing that it engages the stem region (**Fig. 1C**). Consistent with binding the stem, ADI-85666 additionally neutralized a group 2 H7N9 (**Fig. 1B**).

**Fig. 1:**
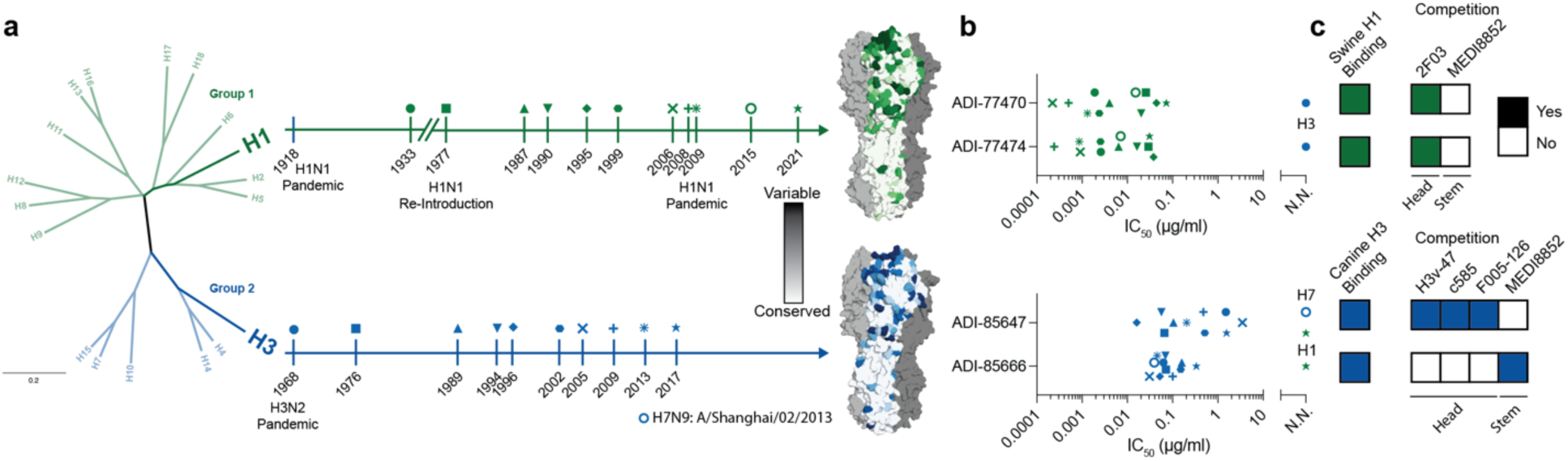
Broad subtype neutralization by human antibodies targeting the HA head. (**A**) Diversity of influenza A HA. Phylogenetic tree of influenza A virus HAs (left). Timeline of influenza H1N1 and H3N2 circulation; symbols represent strains used in this study (middle). Pairwise identity of HAs from viruses used in this study mapped onto structures of HA (PDB: 3LZG and 4FNK) (right). (**B**) Neutralization potencies determined by microneutralization for down-selected antibodies against a panel of influenza A viruses. Strains are indicated as in **A**. H1N1 strains: A/WSN/1933, A/USSR/90/1977, A/Memphis/4/1987, A/Massachusetts/1/1990, A/Beijing/262/1995, A/New Caledonia/20/1999, A/Solomon Islands/3/2006, A/New York/08-1326/2008, A/California/07/2009, A/Michigan/45/2015, A/Sydney/5/2021. H3N2 strains: A/Aichi/2/1968, A/Bilthoven/1761/1976, A/Beijing/353/1989, A/Johannesburg/33/1994, A/Brisbane/8/1996, A/Fujian/411/2002, A/Wisconsin/67/2005, A/Perth/16/2009, A/Switzerland/9715293/2013, A/Kansas/14/2017. (**C**) ELISA binding reactivity at 10 or 1 µg/ml to A/Swine/Henan/SN13/2018^30^ or A/Canine/Hainan/79/2019^31^ (left) and BLI-based antibody competition with the indicated antibodies (right).

### Viral escape from broad H1 antibodies targeting the head

We next more finely mapped the recognition of the broad H1 mAbs by viral escape. We previously generated deep mutational scanning viral libraries of three H1N1s containing all single amino acid mutations and deletions in the HA head (residues 52–263, H3 numbering): A/USSR/90/1977 (USSR77), A/Siena/10/1989 (SN89), and A/Solomon Islands/3/2006 (SI06)^33^. Here, we additionally generated a viral library on the A/Sydney/5/2021 (SYD21) background to determine whether escape is different in a post-2009 pandemic H1N1 virus (**Supplementary Fig. 2A**). We determined neutralization potencies of ADI-77470, ADI-77474, and a V-gene germline-reverted ADI-77474 against these libraries (**Supplementary Fig. 2B**). The germline-reverted ADI-77474 showed undetectable neutralization activity, demonstrating the importance of the somatic mutations in these mAbs (**Supplementary Fig. 2B**). We passaged the four viral libraries in the presence of either ADI-77470 or ADI-77474 at 10x the IC_99_ concentration to identify escape variants (**Fig. 2 and Supplementary Fig. 2B**). Mutation of P122 to acidic residues (e.g., Asp, Glu) or deletion were the most selected in all virus backgrounds (**Fig. 2C and Supplementary Fig. 2F**). However, ADI-77474 was more sensitive to mutational escape in SYD21 compared to its clonal relative ADI-77470 (**Fig. 2 and Supplementary Fig. 2**). Although escape primarily occurred at position 122, mutation to Asn at 122, 123, and 124 in SYD21 were selected by both mAbs; each of these Asn mutations introduces a predicted N-linked glycosylation sequon (**Fig. 2C and Supplementary Fig. 2F**). Although 123N and 124N mutations in the historical strains USSR77, SN89, and SI06 also create potential N-linked glycosylation sites, these mutations were not enriched in these strains, demonstrating different barriers to escape for pre- and post-pandemic H1s (**Fig. 2C and Supplementary Fig. 2F**). We mapped the escape mutations onto the structure of an HA bound by a mAb engaging the same epitope and that uses the same V-genes, V_H_3-23/V_K_1-33^17^ (**Fig. 2B**). Residues 119, 122-125, and 172 cluster around the YxR motif often found in this class of mAbs^17^; the mAbs we describe have a similarly charged YxK motif in the CDRH3 (**Supplementary Fig. 3C**). These escape mutations do not overlap with those of Ab6649^33^, which targets an overlapping region within the lateral patch.

**Fig. 2:**
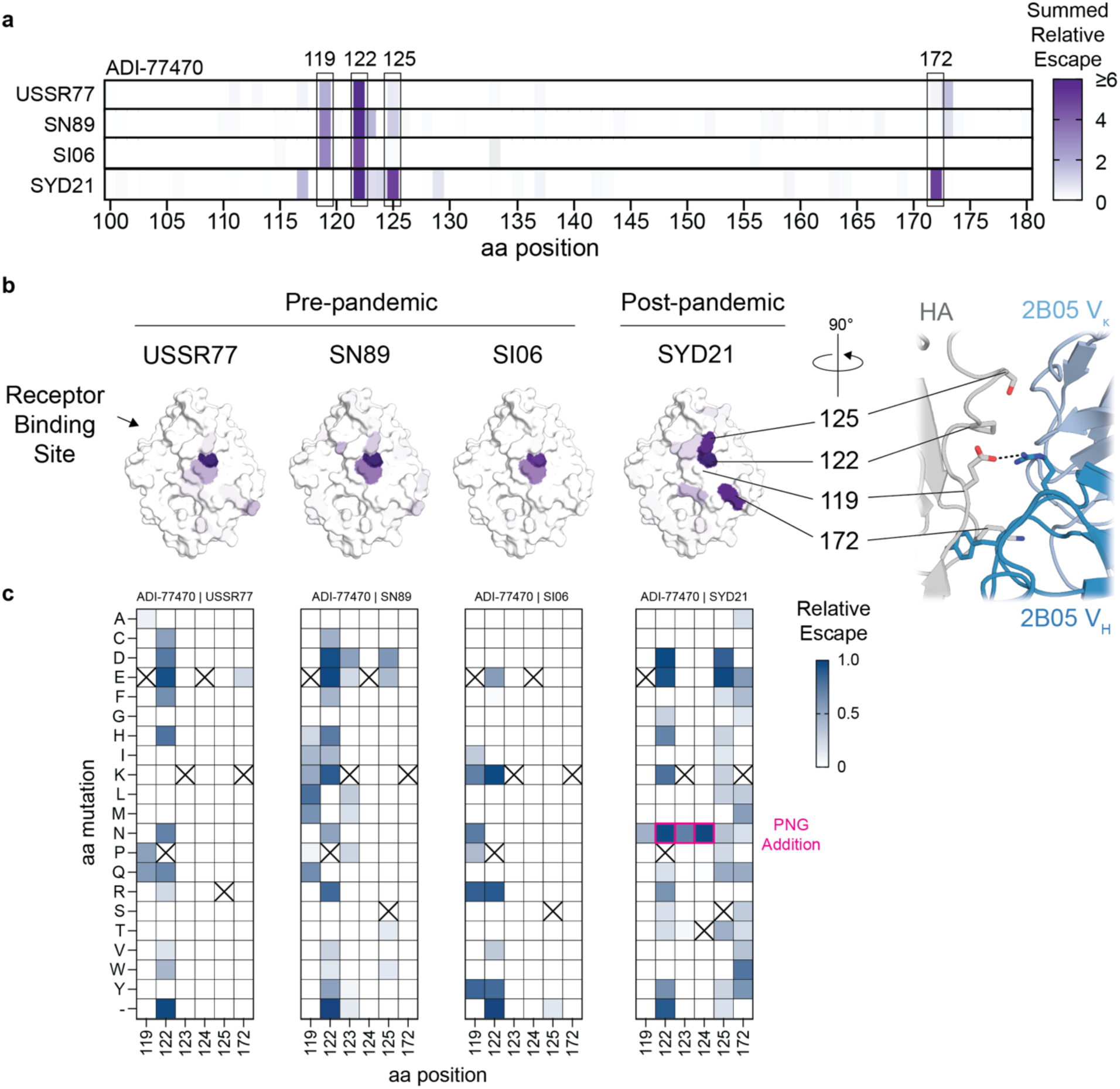
Escape from a broad H1 neutralizing antibody in multiple H1N1 viruses. (**A**) Relative escape values summed by amino acid position in the HA head (H3 numbering) for ADI-77470 against USSR77 (A/USSR/90/1977), SN89 (A/Siena/10/1989), SI06 (A/Solomon Islands/3/2006), and SYD21 (A/Sydney/5/2021). (**B**) Summed relative escape values shown in **A** mapped onto the structure of the SI06 head (PDB: 4HKX) (left) with specific residues highlighted on the structure of the lateral patch-targeting antibody 045-09 2B05 bound to A/California/04/2009 (PDB: 7MEM) (right). (**C**) Relative escape values from ADI-77470 for individual amino acid mutations for each of the strains in **A** for a subset of positions. Highlighted asparagine mutations in SYD21 add potential N-linked glycosylation (PNG) sites.

### Affinity maturation of broad H1 mAbs increases neutralization potency

To increase the affinity of the broad H1 mAbs, we generated yeast libraries encoding mutations in the variable heavy and light chain genes and selected variants with increased binding affinity using recombinant monomeric SYD21 head^34^ (**Supplementary Fig. 3A**). We down-selected affinity-matured mAbs based on a single replicate neutralization screen against SYD21 virus (**Supplementary Fig. 3B**) and further tested the most potent of these mAbs against the historical H1N1 A/Beijing/262/1995 (BE95) (**Fig. 3A**). The *in vitro* affinity-matured mAb, ADI-86325, has 2 heavy and 4 light chain mutations relative to ADI-77470 (**Supplementary Fig. 3C**), which increased the potency against SYD21 by ∼4-fold while maintaining BE95 neutralization (**Fig. 3A**). We next assessed whether *in vitro* affinity maturation of ADI-77470 to ADI-86325 affected viral escape by using deep mutational scanning of the SYD21 virus. We found that mutations that had limited escape effect from ADI-77470 had a reduced effect against ADI-86325, but highly selected escape mutations were nevertheless similar suggesting these residues are critical for HA recognition by these mAbs (**Fig. 2C and 3B**).

**Fig. 3:**
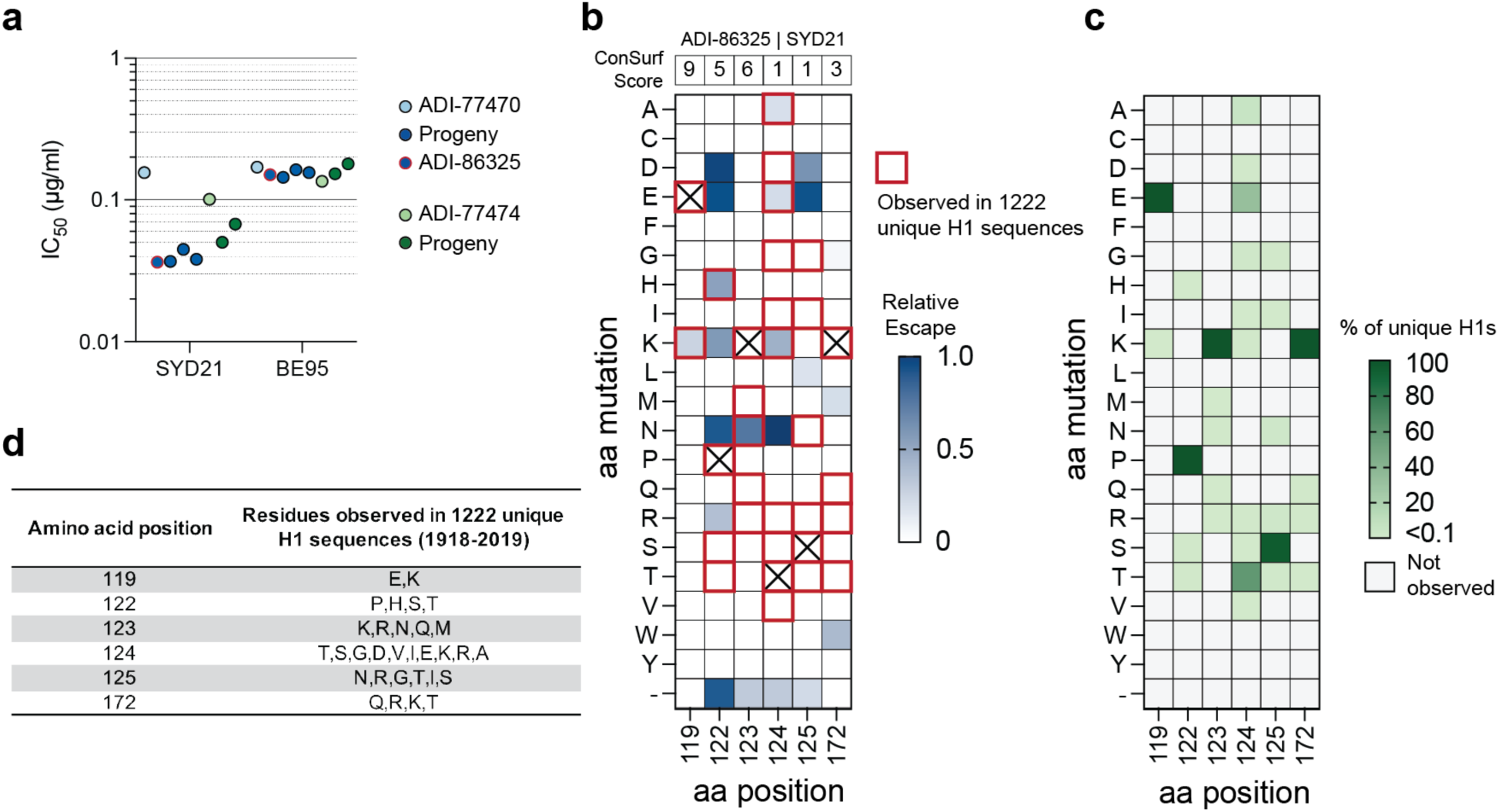
Naturally occurring escape mutations from an engineered lateral patch antibody are rare. (**A**) Neutralization potencies for in vitro affinity-matured antibodies against SYD21 (A/Sydney/5/2021) and BE95 (A/Beijing/262/1995). (**B**) Relative escape values from ADI-86325 for individual amino acid mutations in SYD21 for the positions that can cause escape. ConSurf^36^ values from a previous analysis^35^ are shown at the top. Amino acids boxed in red indicate these were observed in the ConSurf analysis of 1222 H1 sequences that are representative of sequence clusters^35^. (**C**) Frequency of observed mutations in the same set of 1222 unique H1 sequences. (**D**) Table of observed residues at each position.

To determine the occurrence of escape mutations in IAV isolates, we used a previous conservation analysis performed on 1222 unique H1 sequences representative of sequence clusters from 1918–2019^35^ to determine whether observed escape mutations have occurred naturally (**Fig. 3B-D**). Despite some positions with low ConSurf^36^ scores, specific amino acid escape mutations have rarely been observed in H1 sequences (**Fig. 3B-D**), suggesting that ADI-86325 is likely to neutralize most H1N1 viruses and providing an explanation for the high degree of neutralization breadth. The identification of escape mutations *in vitro* that have yet to appear naturally possibly suggests that such antibodies have historically been rare within the human population.

### A broad H3 mAb targets the side of the H3 head

As analogous deep mutational scanning libraries for historical and contemporary H3N2 viruses were unavailable, we used cryo-EM to structurally determine the epitopes targeted by the broad H3 mAbs. ADI-85647, encoded by V_H_3-9/V_K_1-39, in complex with A/Darwin/6/2021 (DAR21) HA shows that it binds to the side of the HA head, consistent with competition with H3v-47, c585, and F005-126 (**Figs. 1, 4A, and Supplementary Fig. 4**). During antigenic drift of H3N2s, potential N-linked glycosylation sites on HA were acquired and maintained at this site at positions 122, 126, and 133^37^. Although the cryo-EM map showed clear features of a glycan at N133, we did not observe a glycan at N122, consistent with a previous report showing no and full glycan occupancy at N122 and N133, respectively^38^. The N133 glycan sits within a cleft between the variable heavy and light chains of ADI-85647, and the CDRH2 inserts between the N133 and N126 glycans (**Fig. 4B**). The Y32 hydroxyl and V2 backbone in the heavy chain form hydrogen bonds with the GlcNAc and one of the branching mannose moieties of the glycan, respectively, and thus stabilize the glycan within this cleft. There are eleven hydrogen bonds, two of which are with the N133 glycan; no salt bridges are present. The heavy chain contributes only two hydrogen bonds in addition to interacting with HA through several aromatic residues (**Fig. 4C**); W102 in the CDRH3 sits within a pocket underneath the 130-loop. While the ADI-85647 epitope significantly overlaps with that of H3v-47 (**Fig. 4D**), ADI-85647 can also bind and neutralize H3N2 strains prior to 1989 in contrast to H3v-47^15^.

**Fig. 4:**
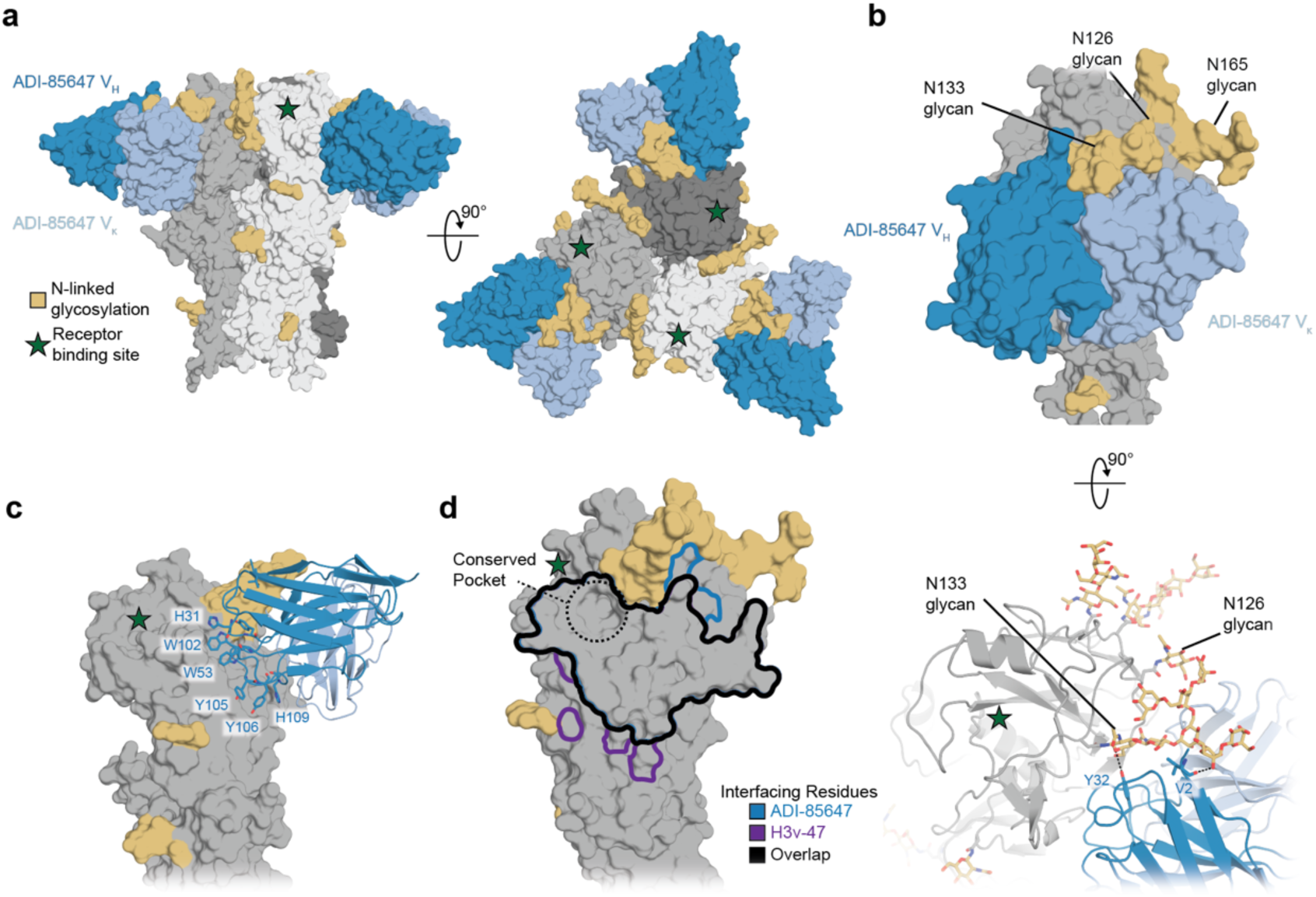
A broad H3 antibody targeting the side of the HA head. (**A**) Cryo-EM structure of ADI-85647 bound to DAR21 (A/Darwin/6/2021) HA. (**B**) A glycan at N133 sits within a cleft formed between the V_H_ and V_κ_ of ADI-85647 (top). Hydrogen bonds formed between the N133 glycan and the V_H_ of ADI-85647 (bottom). (**C**) Several aromatic residues in the heavy chain interact with DAR21 HA. (**D**) The epitopes of ADI-85647 and H3v-47 are highlighted, showing the high degree of overlap between the two epitopes. PISA^53^ was used to determine interfacing residues.

### Group 2 neutralization through a unique structural solution

The anti-stem mAb ADI-85666 uses the germline genes V_H_4-61 and V_K_1-17, which have not been previously reported for group 2 anti-stem mAbs; we therefore determined a cryo-EM structure bound to DAR21 HA_0_ to understand how recognition differs from previously reported mAbs (**Fig. 5A**). The epitope overlaps with recognition of other group 2 neutralizing mAbs^10,39^, such as CR8020, but in contrast to other group 2 neutralizing mAbs, the CDRH3 of ADI-85666 extends into a groove spanning the β-sheet at the base of the stem located near the viral membrane (**Fig. 5B**). The tyrosine at the tip of the CDRH3 makes a stabilizing aromatic-sulfur contact with M133 (HA_2_ numbering) (**Fig. 5B**). One side of the CDRH3 interacts with the hydrophobic groove with I107 and F109 in the CDRH3 while the other side forms a Glu-Arg-Tyr triad with R25 of HA. A second Glu-Arg-Tyr triad is formed with E15 of the fusion peptide and residues Y34 and R99 in the CDRH2 and CDRH3, respectively (**Fig. 5B**). Comparison of the interfacing residues of ADI-85666 with CR8020 shows overlap of the outermost β-strand but diverge towards the central strands or G-helix for ADI-85666 or CR8020, respectively (**Fig. 5C**).

**Fig. 5:**
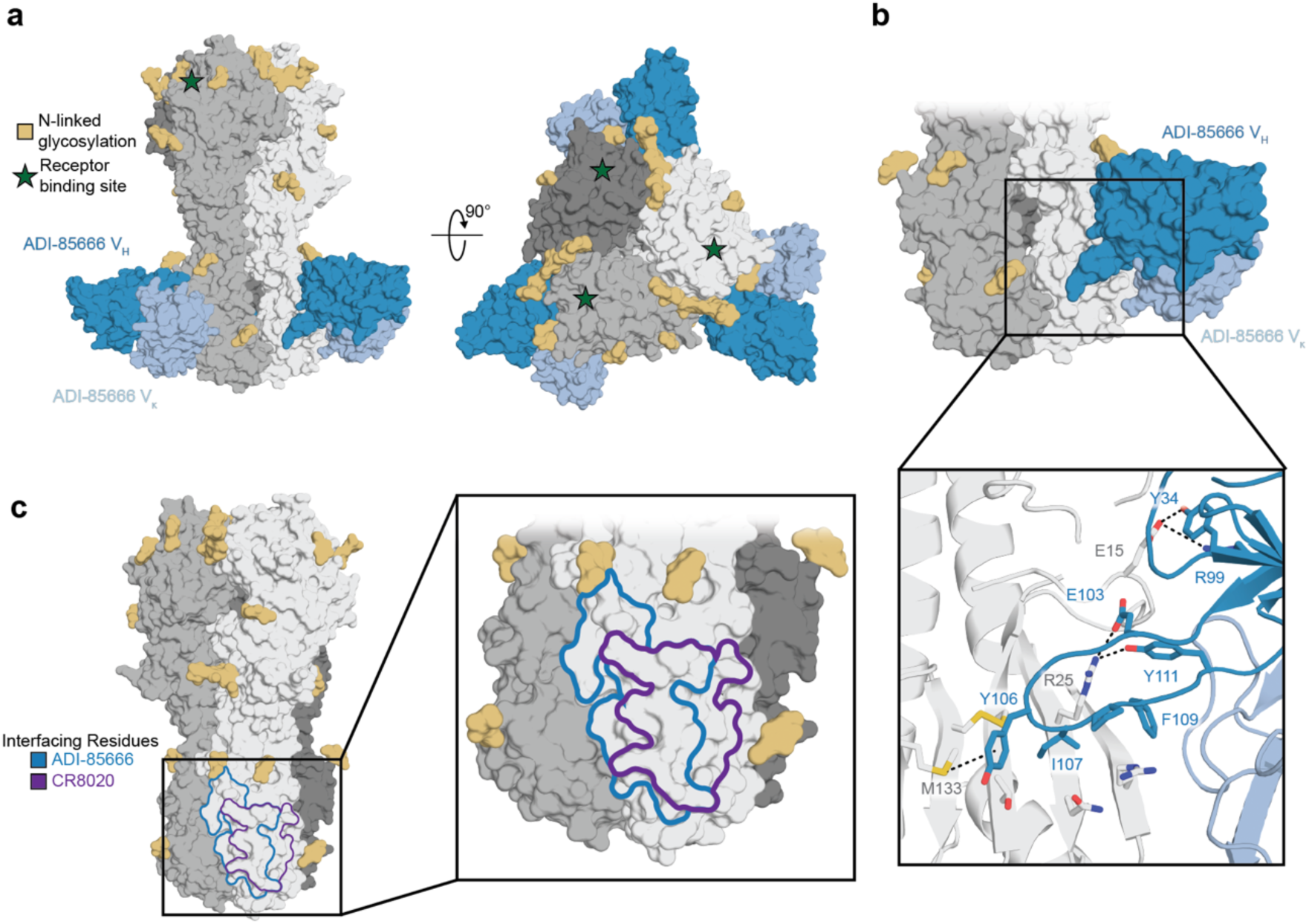
A group 2 neutralizing antibody targets a unique epitope in the stem. (**A**) Cryo-EM structure of ADI-85666 bound to DAR21 (A/Darwin/6/2021) HA. (**B**) Close up of the interactions between the CDRH3 of ADI-85666 and DAR21 HA. (**C**) The epitopes of ADI-85666 and CR8020 are highlighted showing overlapping, but distinct epitopes. PISA^53^ was used to determine interface residues.

### Conservation and mutational tolerance of broad H3 mAb epitopes

We next mapped conservation and mutational tolerance of the ADI-85647 and ADI-85666 epitopes onto the structure of the DAR21 HA (**Fig. 6**). The conservation scores were obtained from a previous ConSurf^36^ analysis of 3,408 unique H3 HA sequences representing historical sequence clusters^35^. To assess the potential for viral escape, we also mapped mutational tolerance derived from A/Perth/16/2009 onto the HA structure, which measures whether mutations at each site impact viral fitness^28^. The ADI-85647 footprint contains some highly variable residues (**Fig. 6**), explaining the variable neutralization potencies (**Fig. 1**); the CDRs of ADI-85647, however, mainly interact with historically conserved residues (**Fig. 6**). Tryptophan 102 in the CDRH3 interacts with a pocket underneath the receptor binding site that is both historically conserved and mutationally intolerant (**Fig. 6**). The groove contacted by ADI-85666 is moderately conserved, but contains few highly variable residues, explaining the broad and consistent neutralization potency (**Fig. 1**). However, several positions within the ADI-85666 footprint are highly tolerant to mutations, suggesting mutation at these sites may result in viral escape (**Fig. 6**).

**Fig. 6:**
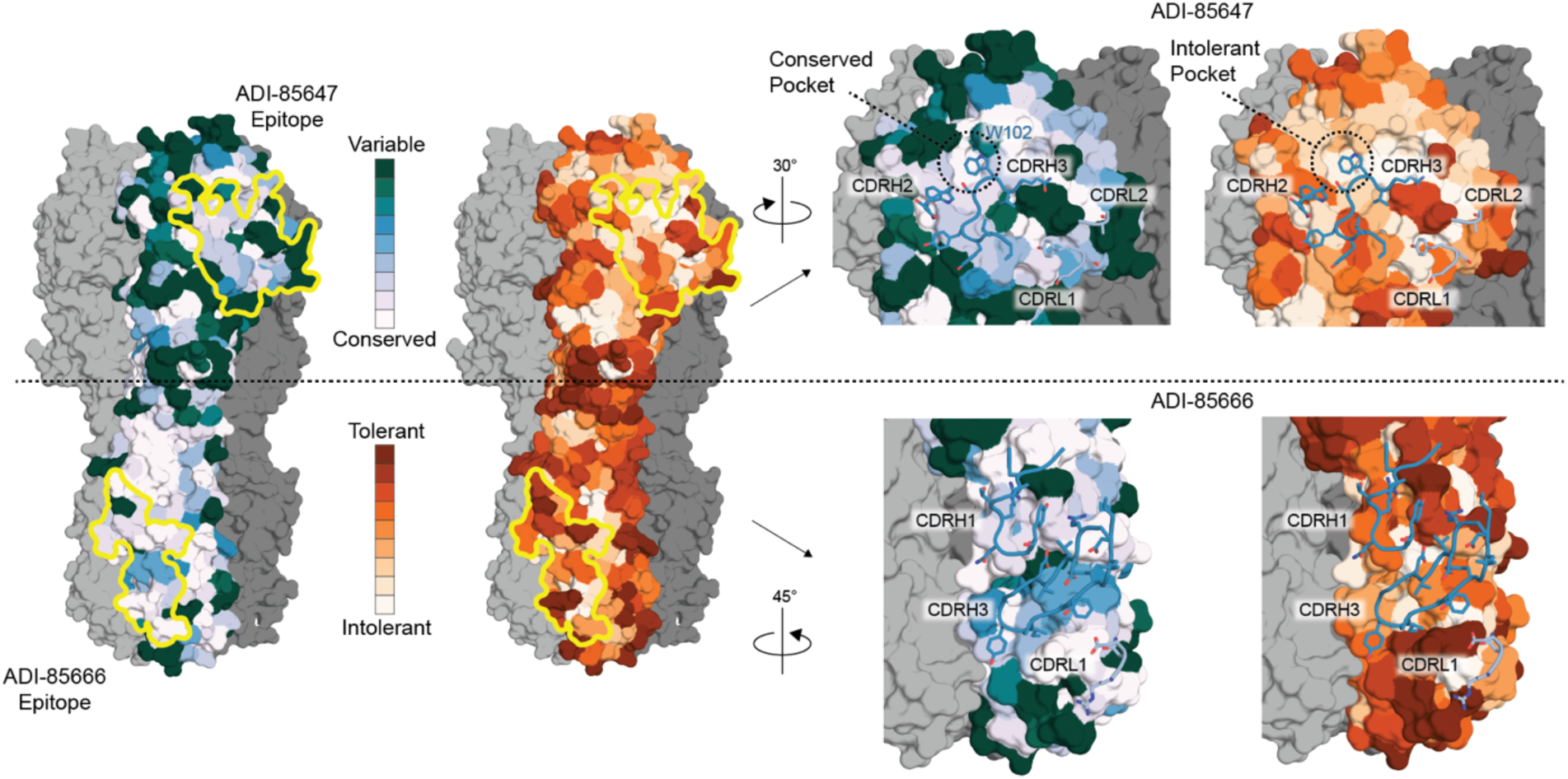
Conservation of epitopes targeted by broad H3 antibodies. ConSurf^36^ scores from 3408 unique H3 HA sequences and mutational tolerance values (0.3-2.7)^28^ are mapped onto the cryo-EM structure of A/Darwin/6/2021 HA. Footprints of ADI-85647 and ADI-85666 are outlined (left). CDRs from ADI-85647 (top, right) and ADI-85666 (bottom, right) are shown in the context of the ConSurf scores and mutational tolerance values. Color spectrums were obtained from colorbrewer2.org.

## DISCUSSION

Here, we isolated broad H1 and H3 neutralizing mAbs from human donors and characterized the targeted epitopes by viral escape and cryo-EM, respectively. Broad H1 mAbs potently neutralize all tested H1N1 strains from 1933 to 2021 and bind to a pre-pandemic swine HA. Despite conservation of the lateral patch recognized by broad H1 mAbs, viral escape occurred readily, primarily at position 122. We engineered the broad H1 mAbs for increased affinity to a contemporary HA, increasing neutralization potency ∼4-fold. This affinity-matured mAb, ADI-86325, selected for viral escape mutations P122D/E/-, 125D/E, and Asn mutations at 122-124, which introduce potential N-linked glycosylation sites; however, these mutations are rarely observed in human IAV isolates. Structural characterization of the broad H3 anti-head mAb, ADI-85647, suggests that engagement of a conserved pocket underneath the 130-loop of the receptor binding site is a target for broad H3 mAbs. The structure of the group 2 anti-stem mAb, ADI-85666, shows the CDRH3 inserts into a groove not contacted by previously characterized group 2 anti-stem mAbs. Comparing these epitopes with previous deep mutational scanning data suggests potential viral escape should this type of antibody response become dominant^28^.

The 2009 H1N1 pandemic initially recalled cross-reactive B cell responses that targeted the RBS and lateral patch^17,22,40^. However, upon subsequent exposures, these responses were absent^17^, which likely contributes to the continued conservation of the lateral patch epitope. Next-generation vaccines using mosaic nanoparticles containing antigenically distinct H1 HA heads elicit antibodies targeting the lateral patch in animal models^25,26^. In humans, initial recall of lateral patch-directed mAbs after the pandemic selected for viral escape through an N-linked glycosylation^22^. However, a subset of lateral patch-directed mAbs, primarily encoded by V_H_3-23/V_K_1-33, can overcome this escape. It is possible that these mosaic nanoparticle vaccine candidates in humans may preferentially elicit V_H_3-23/V_K_1-33 responses targeting the lateral patch. The two clonally related mAbs encoded by V_H_3-23/V_K_1-33 genes we identified here elicited escape at positions engaged by the common CDRH3 motif in this class of mAb (119, 122, 125, and 172), mostly independent of the background strain. Despite further affinity maturation, these mAbs were still readily escaped by mutations at the same positions in contrast to affinity maturation of RBS-directed mAbs that we have previously shown can prevent escape at certain positions^33^. These data suggest that next-generation vaccines eliciting antibodies targeting this site may not provide long-term, broad protection. However, if population-wide immune pressure on this site does not increase, lateral patch mAbs may nevertheless serve as potent prophylactics or therapeutics for high-risk populations. Further studies are needed to understand whether there are mAbs that target this site and are insensitive to the escape mutations described here while maintaining their broad reactivity.

Comparison of the structures of ADI-85647 and H3v-47 show that both insert hydrophobic residues into a conserved pocket underneath the 130-loop. ADI-85647, but not H3v-47, recognizes strains prior to 1989^15^, highlighting that these mAbs are sensitive to different mutations within this epitope despite largely overlapping footprints. As shown previously, the H3 head domain appears to be more mutationally intolerant than the stem; this contrasts with H1 HAs^28^. This conservation and mutational intolerance within the pocket engaged by these mAbs may be a result of the proximity to the conserved W153 in the receptor binding pocket; therefore, mAbs engaging this pocket with different V-genes, angles of approach, and mutation sensitives may be an additional way to target H3N2s with escape resistance; reminiscent of a polyclonal antibody response to the RBS pocket^23^. Although ADI-85647 targets a site on H3 like that targeted by H1 lateral patch mAbs, the neutralization potency is markedly lower. While *in vitro* affinity maturation of ADI-85647 may increase potency, it is possible that H1N1 and H3N2 viruses are differentially neutralized by broad mAbs targeting these sites or that the potency of mAbs targeting these sites are dependent on the angle of approach.

The group 2 mAb, ADI-85666, targets an epitope that overlaps with that of other group 2 neutralizing mAbs, CR8020^10^ and CR8043^39^, but has an extended CDRH3 that reaches into a groove not contacted by other mAbs; this is the third structurally characterized group 2 mAb that engages this site, each with distinct epitopes. The varied gene usage and diversity of recognition suggests that a polyclonal antibody response to this site may be resistant to escape. We note, however, that a recent clinical trial of a prophylactic pan-influenza stem-targeting mAb, VIR2482, failed to significantly reduce IAV illness (NCT05567783). Therefore, although we do not expect group 2 stem-directed mAbs, or vaccines that elicit them, to prevent infection, they may afford an important bridge during seasons of large antigenic change or in the case of spillover of a subtype for which humans have limited immunity. Together, these broad subtypic neutralizing mAbs could be used as prophylactics or therapeutics. Additionally, these mAbs, the viral escape mutations, and structurally characterized epitopes can be used as tools for designing next-generation vaccines for broad protection.

## MATERIALS AND METHODS

### Human B cell sorting

H1 and H3 antigen-specific B cells were detected using recombinant biotinylated H1 and H3 antigens tetramerized with fluorophore-conjugated streptavidin (SA) as previously described^41^. Briefly, His-tagged H1N1 A/California/04/2009 full length trimer Y97F (Sino Biological Cat # 11055-V08B1) was mixed in 4:1 molar ratios with SA-Allophycocyanin (APC; Invitrogen) and SA phycoerythrin (PE; Invitrogen) while A/Darwin/6/2021 full-length soluble trimer and A/Hong Kong/1/1968 head trimer (see below) were mixed in 4:1 molar ratios with SA phycoerythrin (PE; Invitrogen) and SA-Allophycocyanin (APC; Invitrogen), respectively, and were incubated for 20 min on ice. B cells were isolated using a Pan B cell isolation kit (Miltenyi Cat#130-101-638) from approximately 10 million PBMCs and stained with tetramerized antigens (25 nM each); anti-human antibodies anti-CD19 (PE-Cy7; Clone HIB19; Biolegend Cat # 302216), anti-CD3 (PerCP-Cy5.5; CloneOKT3; Biolegend Cat # 317335), anti-CD8 (PerCP-Cy5.5; Clone SK1; Biolegend Cat # 344710), anti-CD14 (PerCP-Cy5.5; Clone 61D3; Invitrogen Cat # 45-0149-42), and anti-CD16 (PerCP-Cy5.5; Clone B73.1; Biolegend Cat # 360712); and 50 μl Brilliant Stain Buffer (BD BioSciences) diluted in FACS buffer (2% BSA/1mM EDTA in 1X PBS). All antibodies were used at 1:100 dilutions and cells were incubated for 15 min on ice. After one wash with FACS buffer, cells were stained in a mixture of propidium iodide and anti-human antibodies anti-IgG (BV605; Clone G18–145; BD Biosciences Cat # 563246), anti-IgA (FITC; Abcam Cat # Ab98553), anti-CD27 (BV510; Clone MT271; BD Biosciences Cat # 740167), and anti-CD71 (APC-Cy7; Clone CY1G4; Biolegend Cat # 334110). Following 15 min of incubation on ice, cells were washed two times with FACS buffer and analyzed using a BD FACS Fusion (BD BioSciences). For sorting of H1/H3-specific, class-switched B cells, B cells that react with H1N1 A/California/04/2009 full length trimer Y97F tetramers or A/Hong Kong/1/1968 head trimer and A/Hong Kong/1/1968 head trimer tetramers among CD19+CD3−CD8− CD14−CD16−PI− and IgG+ or IgA+ cells were single-cell index sorted into 96-well polystyrene microplates (Corning) containing 20 μl lysis buffer per well [5 μl of 5X first strand SSIV cDNA buffer (Invitrogen), 1.25 μl dithiothreitol (0.1 M, Invitrogen), 0.625 μl of NP-40 (Thermo Scientific), 0.25 μl RNaseOUT (Invitrogen), and 12.8 µl dH2O]. Plates were briefly centrifuged and then frozen at −80 °C before PCR amplification.

### Cell lines and media

Cell lines HEK293T, HEK293T-PB1, MDCK-SIAT1-PB1, and MDCK-SIAT1-H7 (gifts from Jesse Bloom and Masaru Kanekiyo) were maintained in DMEM (Gibco) with 10% heat-inactivated FBS (Peak), and penicillin/streptomycin (Gibco). For the growth of MDCK-SIAT1-PB1 and MDCK-SIAT1-H7 cells, 1 mg/ml geneticin (Gibco) and 0.25 µg/ml puromycin (Gibco) were additionally added. For neutralization assays, “flu media” was used, which contains Opti-MEM (Gibco), 0.3% BSA (Sigma), penicillin/streptomycin, 0.1 mg/ml CaCl_2_, and 0.01% heat-inactivated FBS (Peak).

### Generation of influenza reporter viruses

Influenza reporter viruses were generated as described previously^42^ based on the methods from Creanga et al.^35^. Briefly, reverse genetics bidirectional pHW plasmids (some of which were gifts from Jesse Bloom and Masaru Kanekiyo) encoding the HA and NA of each strain, the internal segments (except PB1) from A/WSN/1933 (for H1N1 viruses) or A/Netherlands/009/2010 (for H3N2 viruses), tdKatushka2 with PB1 packing sequences, and a plasmid that drives expression of human TMPRSS2 were transfected into a co-culture of HEK293T-PB1 (a gift from Jesse Bloom) and MDCK-SIAT1-PB1 cells (a gift from Masaru Kanekiyo) in 6-well plates using TransIT-LT1 (MirusBio) at a 2:1 ratio. For H1N1s, the non-coding regions from A/WSN/1933 were used and for H3N2s, the non-coding regions from A/Netherlands/009/2010 were used. The virus-containing supernatant was harvested after three days, clarified by centrifugation, and added to a confluent monolayer of MDCK-SIAT1-PB1 cells seeded the day prior. Once cytopathic effect was evident (two to three days), the supernatant was clarified and stored at −80 °C. To safely generate an H7N9 A/Shanghai/02/2013 virus in a BSL2 setting, we used pHW plasmids encoding tdKatushka with HA packing sequences, the A/Shanghai/02/103 NA, the internal genes from mouse-adapted PR8, and a plasmid that expresses full-length H7 HA from A/Shanghai/02/2013, but lacks non-coding regions. This virus was propagated using the MDCK-SIAT-H7 cell line (a gift from Masaru Kanekiyo) that constitutively expresses cell-surface H7 A/Shanghai/02/2013 HA.

### Microneutralization assays

Neutralization assays were performed as previously reported^43^ based on a previously developed assay^35^. Briefly, MDCK-SIAT1-PB1 (or MDCK-SIAT1-H7 for H7N9 neutralization assays) cells were seeded in 96-well plates (Cellvis, #P96-1.5P) at 20,000-30,000 cells per well in 100 µl of flu media. The next day, viruses were diluted in flu media containing 2 µg/ml TPCK-trypsin and antibodies were diluted separately in flu media at 2-20 µg/ml for screening or titrated to determine IC_50_ values. The virus and antibody dilutions were mixed at a 1:1 ratio (60 µl : 60 µl) and incubated (37 °C / 5% CO_2_) for 1 hour. Then, the flu media was removed from the MDCK-SIAT1-PB1 (or −H7) cells and 100 µl of the antibody/virus mixture was overlayed on top of the cells. Approximately eighteen hours later, the plate was imaged for tdKatushka2 fluorescence using a Zeiss CellDiscoverer7 and fluorescent cells were counted using Zeiss Zen software. The average counts of the virus control wells (no antibody added) determined 0% neutralization and cell only control wells (no virus added) determined 100% neutralization. The neutralization values were plotted in GraphPad Prism and IC_50_ values were determined by a sigmoidal curve fit.

Antibodies used for deep viral escape selections were also titrated on the libraries to determine concentrations to use in selections as described previously^33^. Briefly, 20,000-30,000 humanized MDCK (hCK) cells^44^ were seeded in 96-well plates overnight in flu media. Viruses were then diluted in half-log dilutions for titering or to 200 TCID50 / 50 µl for neutralization assays. For neutralization assays, four-fold antibody dilutions were then added and complexed with virus (1 hour / 37 °C / 5% CO_2_). After 1 hour, the antibody/virus mixture was removed and the cells were washed with PBS before adding fresh flu media with 1 µg/ml TPCK-trypsin. After 18 hours, the cells were fixed with cold 80% acetone in PBS for 10 minutes and air-dried. The wells were washed with PBS + 0.3% Tween20 and blocked with PBS + 0.1% Tween20 + 1% BSA (1 hour, room temperature). The wells were again washed and a mixture of anti-nucleoprotein mAbs (Sigma) were added (1 hour, room temperature). The wells were washed again before adding anti-mouse antibody HRP conjugate (seracare) (1 hour, room temperature). Prior to adding Ultra TMB (ThermoFisher), the wells were washed 5 times. The development of TMB was stopped by addition of 2M sulfuric acid and the resulting absorbance was read at 450 nm. TCID_50_ values were determined by the Reed and Muench method^45^ using 4-8 replicates. Neutralization IC_50_ values were calculated by sigmoidal curve fit in GraphPad Prism and IC_99_ values were then calculated by IC_99_ = (99/(100-99))^(1/hill slope)^ x IC_50_.

### ELISA

Recombinant full-length soluble ectodomain HA (see below) was coated on high binding 96-well plates (Corning) at a concentration of 2 µg/ml overnight at 4 °C. The antigen was then removed and wells were blocked with PBS + 1% BSA + 0.1% Tween20 (1 hour, room temperature). The blocking buffer was removed and 40 µl of antibody was added at 10 or 1 µg/ml to screen for binding (1 hour, room temperature). The antibodies were removed, and the wells were washed three times with PBS + 0.1% Tween20. Anti-human IgG HRP conjugate (Abcam) was then added (1 hour, room temperature). The wells were then washed three times with PBS + 0.1% Tween20 and One Step ABTS was added before reading absorbance at 405 nm. The average of multiple wells with PBS added instead of primary antibody was used to as a background measurement.

### BLI competition assay

Epitope competition binning of mAbs was assessed using the ForteBio Octet HTX (Sartorious). All protein reagents were diluted in PBSF (PBS+0.1% BSA). Anti-human capture (ACH) sensor tips were loaded for 180 seconds with test IgGs at 300 nM. Next, tips were transferred to wells for 180 seconds containing 300 nM soluble HA trimer. After antigen binding, tips were transferred to wells containing test IgGs at 100 nM for 180 seconds to ensure complete blocking of the antigen epitopes of these mAbs. In the last step, sensor tips were transferred to wells containing control IgGs at 100 nM for 180 seconds. An observable binding response in the last step indicates non-competition whereas no observable binding response indicates competition. Data were analyzed using ForteBio Octet BLI analysis software version 12.2.13.5.

### Influenza HA plasmid library generation

Influenza HA library cloning and viral generation was performed as previously described^33^. A deep mutational scanning library encoding all possible single amino acid mutations (Twist Bioscience), including deletions, in the HA head (H3 numbering 52-263) of A/Sydney/5/2021 was assembled using golden gate cloning (BsmBI-v2 Golden Gate Mix, NEB) with a barcoded segment into pDM_RevGen068, a modified pHW2000 containing a GFP insert to detect background colonies during library cloning. After assembly, the reaction was dialyzed prior to electroporation into NEB 10-beta cells. After 1 hour of recovery, the cells were plated on large agar plates and ∼200,000 colonies were scraped, cultured in LB supplemented with carbenicillin for 2 hours before isolating the plasmid library by maxiprep (Macherey-Nagel).

### PacBio sequencing

To link the barcodes with individual amino acid variants, PacBio sequencing of SalI-digested plasmid was performed by the Bauer core using a SMRTbell v3.0 kit and Pacbio Sequel IIe sequencer. Computational methods to link barcodes with amino acid variants was performed exactly as described in Maurer et al.^33^, which is based on previous SARS-CoV-2 RBD deep mutational scanning efforts^46,47^.

### Deep viral escape assay

In T150 flasks with hCK cells (50,000 cells per cm^2^ seeded the day prior in flu media), virus libraries were combined with antibody at 10x the calculated IC_99_ value in 28 mls of flu media with 1 µg/ml TPCK-trypsin and incubated (1 hour / 37 °C / 5% CO_2_). The media was then removed from hCK cells and the virus/antibody mixture was added for 1 hour (37 °C / 5% CO_2_) before removing the inoculum, washing with PBS, and adding fresh flu media with 1 µg/ml TPCK-trypsin. The next day (∼20-24 hours), dead cells and cell debris were removed from the supernatant by centrifugation and 20 mls of clarified supernatant was concentrated with a 100K MWCO concentrator (ThermoFisher). The viral RNA was then extracted with a QIAamp viral RNA mini prep (Qiagen).

### Viral barcode amplification

Barcode amplification was performed as previously described^33^. One µl ezDNase buffer and 1 µl ezDNase was added to 8 µl of viral RNA and incubated at 37 °C for 2 minutes. Then, 1 µl dNTPs (10 mM), 1 µl water, and a primer specific for the 3’ NCR (prDM370) was added. The mixture was incubated for 5 minutes at 65 °C followed by ice for 1 minute. To this mixture, 1 µl SuperScript IV, 4 µl SuperScript IV buffer, 1 µl RNaseOUT, and 1 µl DTT were added and the reaction was incubated at 50 °C for 30 minutes and then 80 °C for 10 minutes. The cDNA was purified by a 1.6X AMPure XP bead cleanup (Beckman). The cDNA was amplified in two PCR steps. The first PCR step combined 20 µl cDNA, 10 µl Q5 buffer, 1 µl dNTPs, 13.5 µl water, 2.5 µl each primer, and 0.5 µl Q5 polymerase. The cycling conditions for the first PCR step were 98 °C for 30 seconds; four cycles of 98 °C for 10s, 67 °C for 10s, 72 °C for 10s; 72 °C for 2 minutes. The first PCR was purified by a 1.6X AMPure XP bead cleanup. The second PCR step combined 20 µl PCR1 product, 10 µl Q5 buffer, 1 µl dNTPs, 13.5 µl water, 5 µl NEBNext unique dual index primers, and 0.5 µl Q5 polymerase. The cycling conditions for the second PCR step were 98 °C for 30s; 14 cycles of 98 °C for 10s, 72 °C for 15s; 72 °C for 2 minutes. After purification with AMPure XP beads, the product was analyzed by a D1000 tapestation, and combined at equimolar concentrations before quantifying with Qubit and submitting for sequencing on a NovaSeq SP (Bauer Core).

### Protein production

Antibodies used in selections and recombinant HAs (head or full-length soluble ectodomain) were cloned into a pVRC8400 (Vaccine Research Center, NIAID, Bethesda, MD) backbone. All proteins contained a His-tag and were purified by cobalt chromatography followed by FPLC purification on S200 column using an AKTA Pure. All proteins, except for the A/Darwin/6/2021 HA trimer, were produced in HEK293F by transfecting with PEI (3:1 ratio) or Expi-293F (ThermoFisher) cells following the manufacturer’s recommended transfection protocol. Full-length soluble ectodomain trimers, except for the A/Darwin/6/2021 HA trimer used in cryo-EM, contained a foldon trimerization tag. The A/Darwin/6/2021 HA trimer used for cryo-EM was produced in Expi293F GnTI-cells and contained three stabilizing mutations in the stem: H26W, K51I, and E103I^48^. Proteins used for B cell sorting and yeast-based affinity maturation were biotinylated by amine coupling and quantified using the QuantTag biotin kit (Vector Laboratories, #BDK-200). Antibodies used for screening, as well as ADI-86325 used in deep mutational scanning selections, were produced in yeast. Expression and purification of IgGs in yeast were produced as full-length IgG1 proteins in S. cerevisiae cultures, as previously described^49^. Briefly, yeast cultures were incubated in 24-well plates placed in Infors Multitron shaking incubators at 30 °C, 650 rpm, and 80% relative humidity. After 6 days, the supernatants containing the IgGs were harvested by centrifugation and purified by protein A-affinity chromatography. The bound IgGs were eluted with 200 mM acetic acid with 50 mM NaCl (pH 3.5) into 1/8 [v/v] 2 M HEPES (pH 8.0) and buffer-exchanged into PBS (pH 7.0).

### Yeast-based affinity maturation

Affinity maturation libraries were generated as described previously^50^. Overlap extension PCR was used to incorporate NNK codons into the complementary determining regions (CDRs) of the variable heavy- and light-chains. Variable heavy and light chain libraries were generated separately and recombined in heavy chain or light chain libraries in yeast by homologous recombination with a linearized plasmid to create yeast libraries of ∼1 x 10^7^ by electroporation. To prepare S. cerevisiae yeast for electroporation, cells were first grown overnight in YPD media and refreshed the morning of electroporation to 0.5 OD600 in YPD for 4 hours. Yeast cells were then collected and washed twice with sterile water before incubating in 0.1 M lithium acetate / 10 mM DTT for 10 minutes. The yeast cells were then washed twice with ice-cold water and resuspended in ice-cold 10 mM lithium acetate. The insert and plasmid DNA were added to the yeast cells and electroporated at 0.5 kV / 2000 µF. YPD media was used to recover the cells before incubating at 30 °C. The yeast cells were then resuspended in SDCAA media (6.7 g/L Yeast Nitrogen Base, 4.0 g/L drop out amino acid mix, 7.3 g/L NaH2PO4, 5.4 g/L Na2HPO4, 4% w/v dextrose) and grown at 30 °C. The heavy- and light-chain libraries were selected over two rounds for binding to recombinant A/Sydney/5/2021 HA head monomer. The number of yeast cells stained in each selection was at least 10-fold the library diversity. Each library was incubated with either 10 or 1 nM of biotinylated SYD21 HA head monomer under equilibrium conditions. The yeast cells were washed twice in PBSF (1x PBS, 0.1% [w/v] BSA) before staining with anti-human LC-FITC (Southern Biotech), Streptavidin 633 (Invitrogen), and propidium iodide (Invitrogen) for 15 minutes on ice. The yeast cells were washed twice and resuspended in PBSF before sorting on a BD FACS Aria II. Yeast cells showing improved antigen binding over parental clones were sorted in each round. After two rounds of sorting, the outputs from the heavy- and light-chain libraries were combined to generate a library of clone with diversity across all CDRs. This library was also sorted for two rounds before plating the yeast on SDCAA agar plates for single clone isolation and sequencing.

### HA conservation and mutational tolerance analyses

For the pairwise identity maps shown in Figure 1, HA sequences of reporter viruses used in this study were aligned and pairwise identity was calculated in Geneious. For conservation analyses of H1 and H3 HAs we used a previous analysis^35^ of 1222 unique H1 or 3408 unique H3 HA sequences from the GISAID database^51^, removing duplicates or sequences with poor sequencing data, performing clustering using CD-HIT EST^52^, and selecting representative sequences from these clusters. Cluster representatives were then analyzed for conservation using ConSurf^36^. For mutational tolerance analyses, we used the entropy values from Lee et al.^28^. Color spectrums were obtained from colorbrewer2.org. Antibody footprints were determined by interfacing residues from PISA^53^.

### Cryo-EM sample preparation and data collection

The recombinant HA (rHA) full length soluble ectodomain (FLsE) of A/Darwin/6/2021 (H3N2) was incubated with ADI-85647 or ADI-85666 Fab at a molar ratio 1:1 and a final protein concentration of 2.5 mg/ml for 40 minutes on ice. Immediately before depositing the sample onto Quantifoil R0.6/1, 3400 mesh gold grids coated with 2nm continuous carbon, octyl-beta-glucoside was added to the pre-formed complex at 0.083% (w/v) final concentration. Plasma cleaning of the grids was performed with a Solarus plasma cleaner machine (Gatan) for 30 seconds (H2, O2). Three microliters of the sample were applied on the continuous carbon face of the grids, blotted for 5.5 seconds on the same face of the grid, at 20° C and 80% humidity, then plunge-frozen in 100% liquid ethane using a Leica EM GP2 plunge freezer. The grids were first screened on an Artica Talos microscope equipped with a Gatan K3 camera and operated at 200 kV. The highest quality grid was then imaged using a Titan Krios microscope operated at 300 kV and equipped with a Gatan K3 Summit direct electron detector. A total of 7,772 and 4,435 movies for the ADI-85647 or ADI-85666 complex, respectively, were collected in counting mode at 30 e-/px/s for a total dose of 52.44 e^−^/Å^2^/s. Images were collected at a magnification of 81000, corresponding to a calibrated pixel size of 0.825 Å/pixel. Defocus values ranged between −1 and −2.5 µm.

### Cryo-EM data processing

The data were processed (Supplementary Figs. 5 and 6) using cryoSPARC v4.3.1^54^. Briefly, the movies were aligned and corrected for both global and local motion using Patch Motion Correction within cryoSPARC. Then, contrast transfer function estimation was done using Patch CTF and particles were picked using template picker. The picked particles were extracted with a box size 400 pixels and 4x binned for 2D classification. Two rounds of 2D classification were performed and the good particles were selected for the generation of the initial model at 12 Å/pixel with 3 classes. Non-uniform refinement was performed in C1 symmetry for each individual class. The particles from the best class were re-extracted with a box size of 400 pixels and binned 2x and then subject to 3D classification into 3 classes with a target resolution of 4Å. The particles from the top class were combined and used for non-uniform refinement and local CTF refinement with the best volume obtained from the 3D classification. The particle stack from the local CTF refinement was then used for new round of non-uniform refinement, and then a new round of 3D classification was performed with 3 classes at a target resolution of 3Å. The 3 classes were subject to a new round of non-uniform refinement. Local CTF refinement was then performed the final particle stack and then subjected to a new round of non-uniform refinement with C3 symmetry imposed. A new round of local CTF refinement followed by non-uniform refinement in C3 symmetry was performed and then subject to global CTF refinement follow by new round of non-uniform refinement in C3 symmetry. The particles were re-extracted at full box size of 400 pixels without binning and a new round of non-uniform refinement and local CTF refinement was performed. This particle stack was used for a last non-uniform refinement with C3 symmetry and yielded a reconstruction at 3Å nominal resolution. The two half-maps were used for sharpening in DeepEMhancer^55^. The reported resolution is based on the gold-standard Fourier shell correlation of 0.143 criterion.

### Model building and refinement

The DeepEMhancer sharpened maps were used for model building with ModelAngelo^56^ and then manually built using COOT^57^. N-linked glycans were built manually in COOT using the glyco extension and their stereochemistry and fit to the map validated with Privateer^58^. The models were then refined in Phenix^59^ using real-space refinement and validated with MolProbity^60^.

### Computational sequencing analyses

Computational sequencing analyses were performed exactly as described in Maurer et al.^33^. Briefly, barcodes and amino acid variants were linked from PacBio sequencing using *alignparse*^61^. Variants with insertions were removed from analysis. Sequences with ≥3 PacBio CCSs per barcode were retained for analysis. After amplification of the barcodes, paired ends were merged with PEAR^62^, merged reads were filtered for the correct lengths, and sequences with ambiguous base calls were removed. Unique barcode-UMI combinations were used for counting. The functional scores were calculated using *dms_variants* (https://jbloomlab.github.io/dms_variants/) comparing the change in variant frequency from the plasmid libraries to after the low MOI (0.01) passage. Relative escape values are derived from differential selection (https://jbloomlab.github.io/dms_tools2/diffsel.html) using a pseudocount of 10. For analysis, we filtered for variants that were positively selected in both independent replicates and set other variants to 0. Relative escape values are normalized to the most selected variant in each selection. These relative escape values are summed by position to obtain a “summed relative escape” value.

### Data and code availability

Raw sequencing data have been deposited to NCBI in BioProjects PRJNA1130551 and PRJNA1056469. All code used for the analyses here and the resulting output files used to generate the figures can be found at https://github.com/dpmaurer/flu_ab_escape. Atomic models have been deposited to the PDB under accession codes 9BDF and 9BDG and the corresponding maps have been deposited to the EMDB under accession codes EMD-44451 and EMD-44452. Aligned micrographs for these datasets can be found in EMPIAR under accession codes EMPIAR-12059 and EMPIAR-12060.

## ACKNOWLEDGMENTS

At Adimab, we thank the sequencing and yeast antibody production teams, as well as Irina Burina and Elizabeth McGurk for BLI competition assays. We thank Jesse Bloom for providing pHW plasmids. We thank Masaru Kanekiyo for providing the HA and NA plasmids used to generate the reporter viruses and for the cell lines used to propagate the reporter viruses. We thank Adrian Creanga for providing the ConSurf analyses. We acknowledge support from P01 AI089618 (A.G.S). This research has been funded in whole or part with federal funds under a contract from the National Institute of Allergy and Infectious Diseases, NIH contract 75N93019C00050 (A.G.S. and G.B.). We also acknowledge support from the Irma T. Hirschl/Monique Weill-Caulier Trust (G.B.). Some of this work was performed at the National Center for CryoEM Access and Training (NCCAT) and the Simons Electron Microscopy Center located at the New York Structural Biology Center, supported by the NIH Common Fund Transformative High Resolution Cryo-Electron Microscopy program (U24 GM129539), and by grants from the Simons Foundation (SF349247) and NY State Assembly.

## AUTHOR INFORMATION

### Author Contributions

D.P.M. and M.V. produced recombinant proteins, performed ELISAs, and performed virus neutralization assays. D.P.M. performed deep mutational scanning assays and analyses. H.L.D., P.K., and J.C.G. sorted human B cells and performed *in vitro* affinity maturation. A.S.F.R. and G.B. determined cryo-EM structures. D.P.M. and A.G.S. wrote the original drafts of the paper. All authors reviewed and commented on the paper.

**Correspondence and requests for materials should be addressed to:** Aaron G. Schmidt

