## Supplementary Materials for "Conserved sites on the influenza H1 and H3 hemagglutinin recognized by human antibodies"

[*]These authors contributed equally

### Supplementary Table 1: Cryo-EM data collection and statistics.

| | H3/Darwin/6/2021 in complex with ADI-85647 | H3/Darwin/6/2021 in complex with ADI-85666 |
|---|---|---|
| | PDB-9BDG, EMD-44452 and EMPIAR-12059 (deposition ID: 47485795) | PDB-9BDF, EMD-44451 and EMPIAR-12060 (deposition ID: 47485853) |
| | **Data Collection** | |
| **Grid type** | 2nm UTC Quantifoil 0.6/1.0 300 | 2nm UTC Quantifoil 0.6/1.0 300 |
| **Microscope/voltage/detector** | Titan Krios/300 kV/Gatan K3 summit | Titan Krios/300 kV/Gatan K3 summit |
| **Energy filter** | 20 eV slit | 20 eV slit |
| **Magnification** | 81,000 | 81,000 |
| **Recording mode** | counting | counting |
| **Total dose** | 52.44 e-/Å$^2$/s | 52.44 e-/Å$^2$/s |
| **Pixel size** | 0.825 Å/pixel | 0.825 Å/pixel |
| **Defocus range** | –0.8 to –2 µm | –0.8 to –2 µm |
| **No. micrographs used** | 7,772 | 4,435 |
| **Total particles picked** | 4,789,730 | 3,009,387 |
| | **Model Validation** | |
| **Composition (#)** | | |
| **Chains** | 9 | 9 |
| **Atoms** | 15200 | 17167 |
| **Residues** | Protein: 1853 Nucleotide: 0 | Protein: 2126 Nucleotide: 0 |
| **Ligands** | BMA: 9; NAG: 35; MAN: 21 | BMA: 4; NAG: 34; MAN: 3 |
| | **Bonds (RMSD)** | |
| **Length (Å) (# > 4sigma)** | 0.004 (0) | 0.003 (0) |
| **Angles (°) (# > 4sigma)** | 0.614 (0) | 0.583 (0) |
| **MolProbity score** | 1.88 | 1.82 |
| **Clash score** | 6.27 | 4.82 |
| | **Ramachandran plot (%)** | |
| **Outliers** | 0 | 0 |
| **Allowed** | 4.92 | 3.66 |
| **Favored** | 95.08 | 96.34 |
| **Rotamer outliers (%)** | 1.81 | 2.66 |
| **Cbeta outliers (%)** | NA | NA |
| | **Peptide plane (%)** | |
| **Cis proline/general** | 2.4/0.0 | 0.0/0.0 |
| **Twisted proline/general** | 0.0/0.0 | 0.0/0.0 |
| **CaBLAM outliers (%)** | 2.27 | 2.26 |
| | **ADP (B-factors) min/max/mean** | |
| **Protein** | 25.15/172.01/68.96 | 25.15/172.01/68.96 |
| **Ligand** | 65.65/110.11/87.89 | 65.65/110.11/87.89 |
| | **Data** | |
| **Lengths (Å)** | 146.03, 150.15, 124.58 | 142.72, 136.95, 145.20 |
| **Angles (°)** | 90.00, 90.00, 90.00 | 90.00, 90.00, 90.00 |
| **Supplied Resolution (Å)** | 3.1 | 3 |
| **Resolution Estimates (Å)** | Masked | Masked |
| **d FSC (half maps; 0.143)** | --- | --- |
| **d 99 (full/half1/half2)** | 2.3/---/--- | 2.3/---/--- |
| **d model** | 2.1 | 2.1 |
| **d FSC model (0/0.143/0.5)** | 1.6/1.7/3.0 | 1.5/1.7/3.0 |
| **Map min/max/mean** | 0 | 0 |
| | **Model vs. Data** | |
| **CC (mask)** | 0.85 | 0.85 |
| **CC (box)** | 0.77 | 0.79 |
| **CC (peaks)** | 0.72 | 0.76 |
| **CC (volume)** | 0.85 | 0.86 |
| **Mean CC for ligands** | 0.69 | 0.74 |

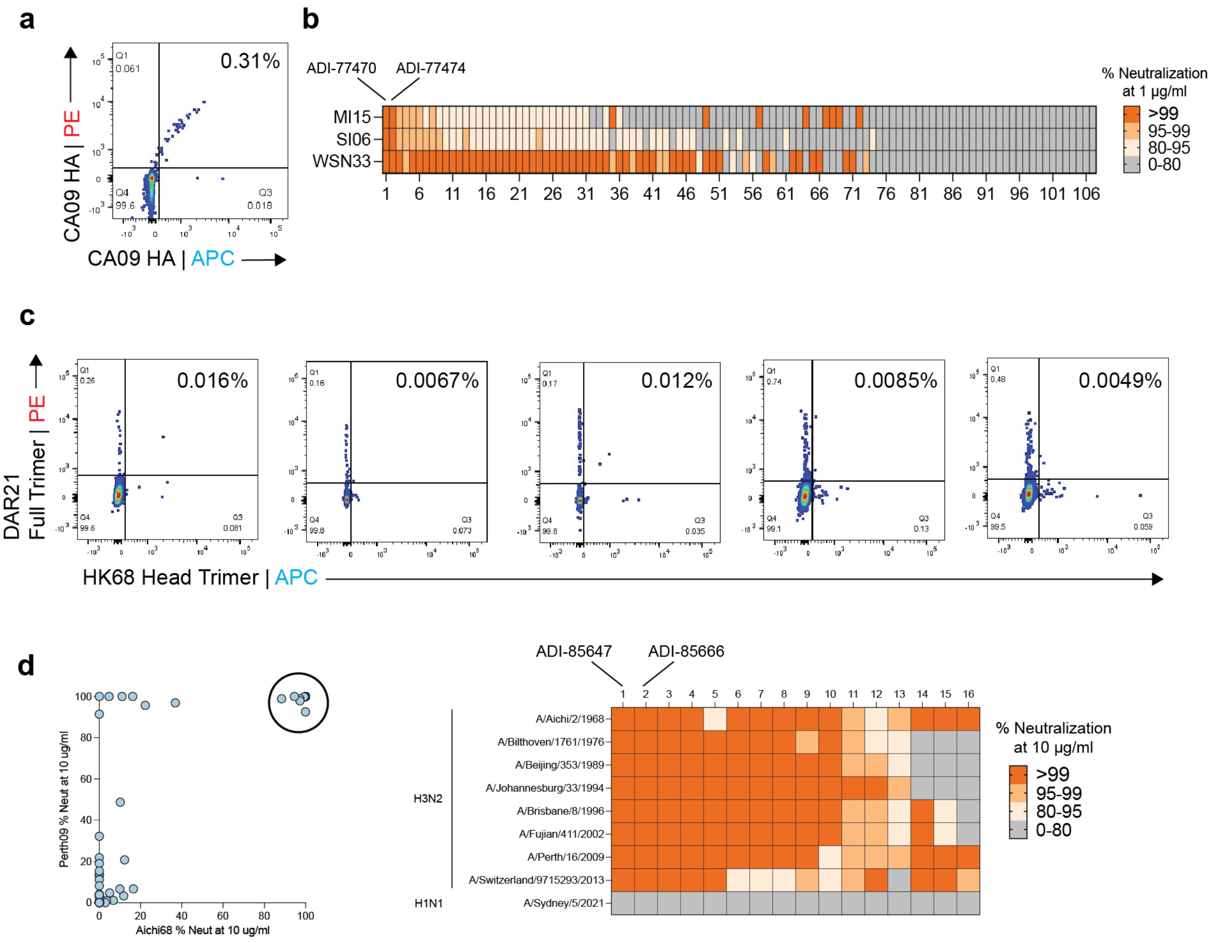


**Supplementary Fig. 1: B cell sorting and antibody screening.** (**A**) Sort plot gated on CD19$^+$ and IgA/IgG$^+$ B cells to isolate A/California/07/2009 HA-reactive B cells. (**B**) Neutralization screening at 1 µg/ml of recombinant antibodies against A/Michigan/45/2015, A/Solomon Islands/3/2006, and A/WSN/1933. (**C**) Sort plots for individual donors gated on CD19$^+$ and IgA/IgG$^+$ B cells to isolate B cells cross-reactive between A/Darwin/6/2021 full-length soluble trimer and A/Hong Kong/1/1968 head trimer. (**D**) Cross-neutralization screen between A/Aichi/2/1968 and A/Perth/16/2009 (left). Circled clones (n=16) were screened at 10 µg/ml against a panel of H3N2 reporter viruses (right).

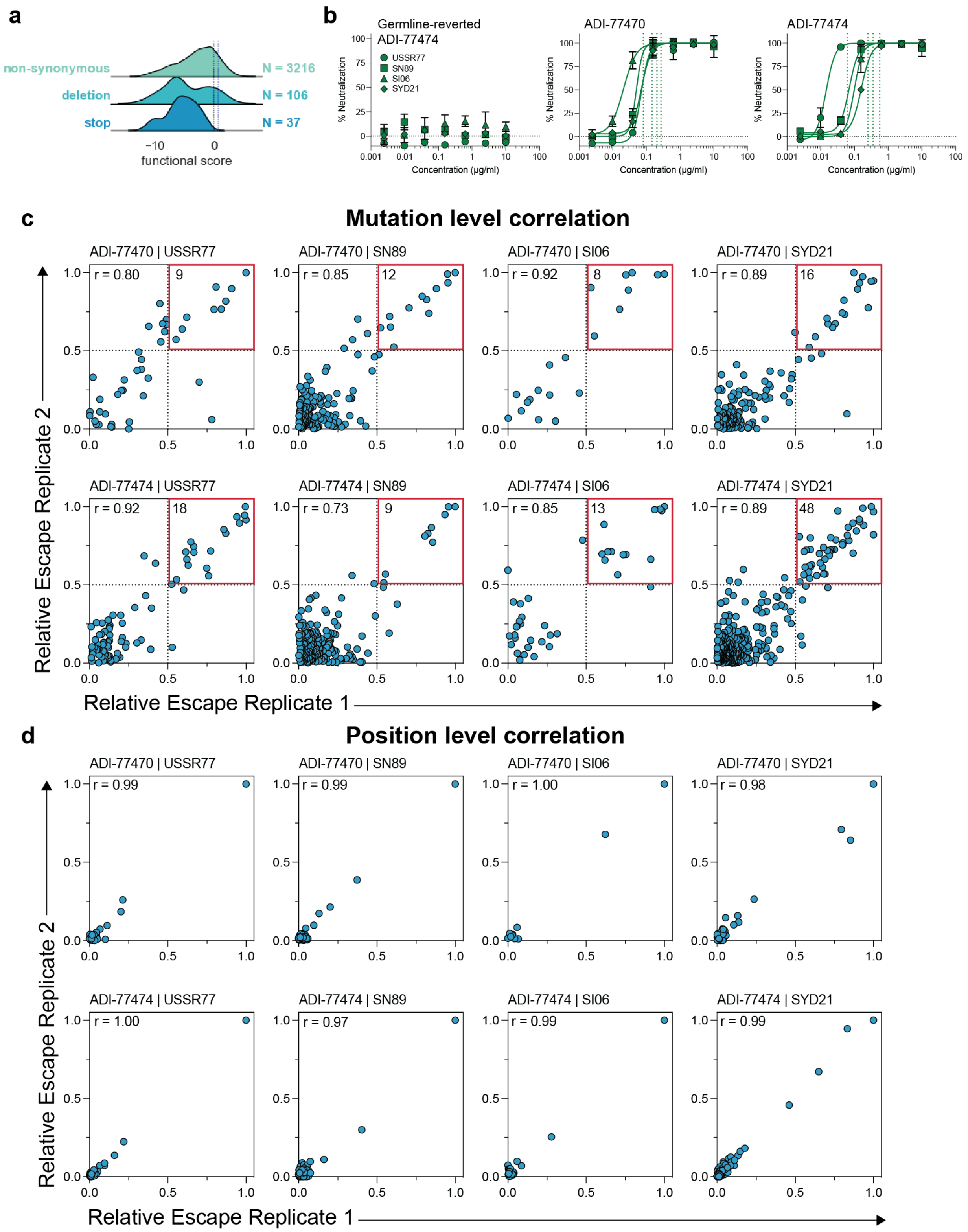
a
non-synonymous
N = 3216
deletion
N = 106
stop
N = 37
−10
0
functional score
b
Germline-reverted ADI-77474
USSR77
SN89
SI06
SYD21
% Neutralization
Concentration (μg/ml)
ADI-77470
ADI-77474
c
Mutation level correlation
ADI-77470 | USSR77
r = 0.80
9
ADI-77470 | SN89
r = 0.85
12
ADI-77470 | SI06
r = 0.92
8
ADI-77470 | SYD21
r = 0.89
16
ADI-77474 | USSR77
r = 0.92
18
ADI-77474 | SN89
r = 0.73
9
ADI-77474 | SI06
r = 0.85
13
ADI-77474 | SYD21
r = 0.89
48
Relative Escape Replicate 2
Relative Escape Replicate 1
d
Position level correlation
ADI-77470 | USSR77
r = 0.99
ADI-77470 | SN89
r = 0.99
ADI-77470 | SI06
r = 1.00
ADI-77470 | SYD21
r = 0.98
ADI-77474 | USSR77
r = 1.00
ADI-77474 | SN89
r = 0.97
ADI-77474 | SI06
r = 0.99
ADI-77474 | SYD21
r = 0.99
Relative Escape Replicate 2
Relative Escape Replicate 1

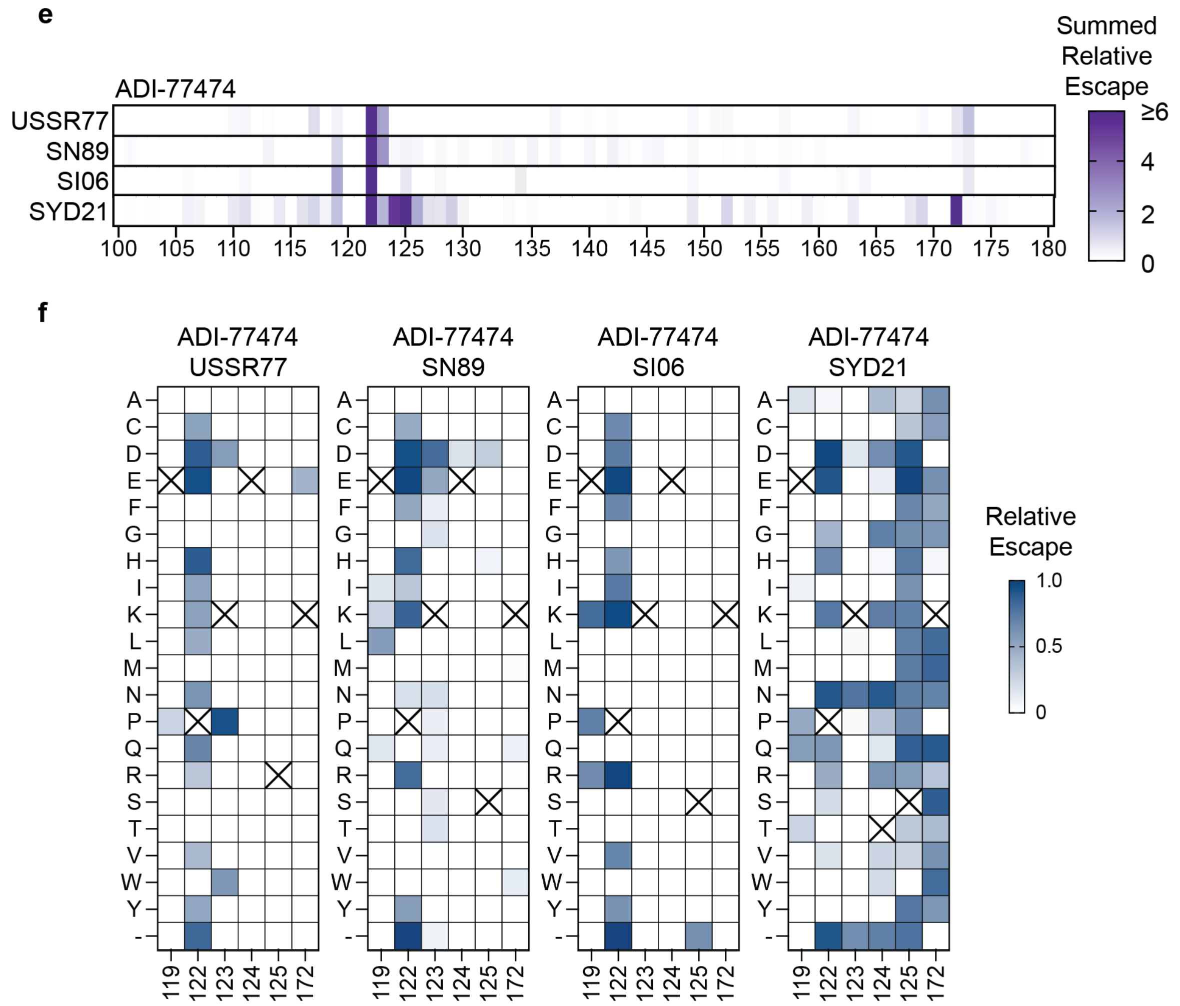


**Supplementary Fig. 2: Deep mutational scanning selections.** (**A**) Functional scores determined after a low MOI (0.01) passage for non-synonymous, deletion, and stop mutations. Black dotted line is the functional score for wild-type A/Sydney/5/2021 and the blue dotted line is the functional score of synonymous mutants within the library. (**B**) Neutralization of the four deep mutational scanning libraries with the indicated antibody. (**C**) Correlation between independently generated replicate libraries by individual amino acid mutations for each selection. (**D**) Correlation between independently generated replicate libraries summed by amino acid position for each selection. (**E**) Relative escape values from ADI-77474 summed by amino acid position in the HA head of the indicated viruses: A/USSR/90/1977, A/Siena/10/1989, A/Solomon Islands/3/2006, and A/Sydney/5/2021. (**F**) Relative escape values from ADI-77474 of individual amino acid mutations at highly selected positions.

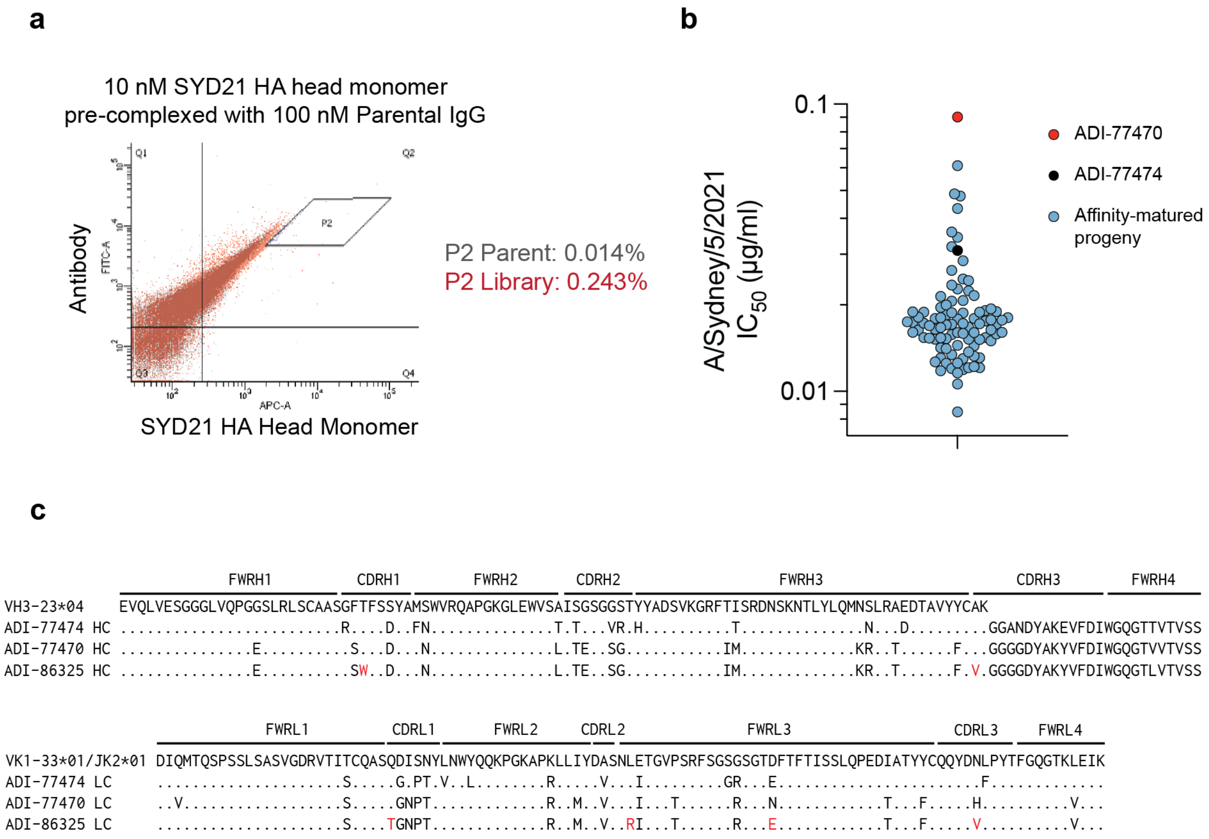


**Supplementary Fig. 3: In vitro affinity maturation of ADI-77470 and ADI-77474.** (**A**) Terminal sorting round with A/Sydney/5/2021 (SYD21) HA head pre-complexed with parental IgG. (**B**) Neutralization screening of affinity-matured progeny. (**C**) Sequences of ADI-77470, ADI-77474, and the top affinity-matured clone ADI-86325 relative to germline.

```
                          FWRH1             CDRH1        FWRH2       CDRH2                  FWRH3                       CDRH3          FWRH4
VH3-9*01      EVQLVESGGGLVQPGRSLRLSCAASGFTFDDYAMHWVRQAPGKGLEWVSGISWNSGSIGYADSVKGRFTISRDNAKNSLYLQMNSLRAEDTALYYCAK
ADI-85647 HC  ............................HN...............D..T..T...VNVD.....................................V.GQVWSGYYLTHDAFDLWGQGTTVTVSS
```

```
                          FWRL1           CDRL1      FWRL2      CDRL2              FWRL3               CDRL3   FWRL4
VK1-39*01     DIQMTQSPSSLSASVGDRVTITCRASQSISSYLNWYQQKPGKAPKLLIYAASSLQSGVPSRFSGSGSGTDFTLTISSLQPEDFATYYCQQSYSTP
ADI-85647 LC  ..V.....P....T................N.................SG.....L.........E................C.......G.IM.GTFGQGTKVDIK
```

**Supplementary Fig. 4: Sequence of ADI-85647 relative to germline.**

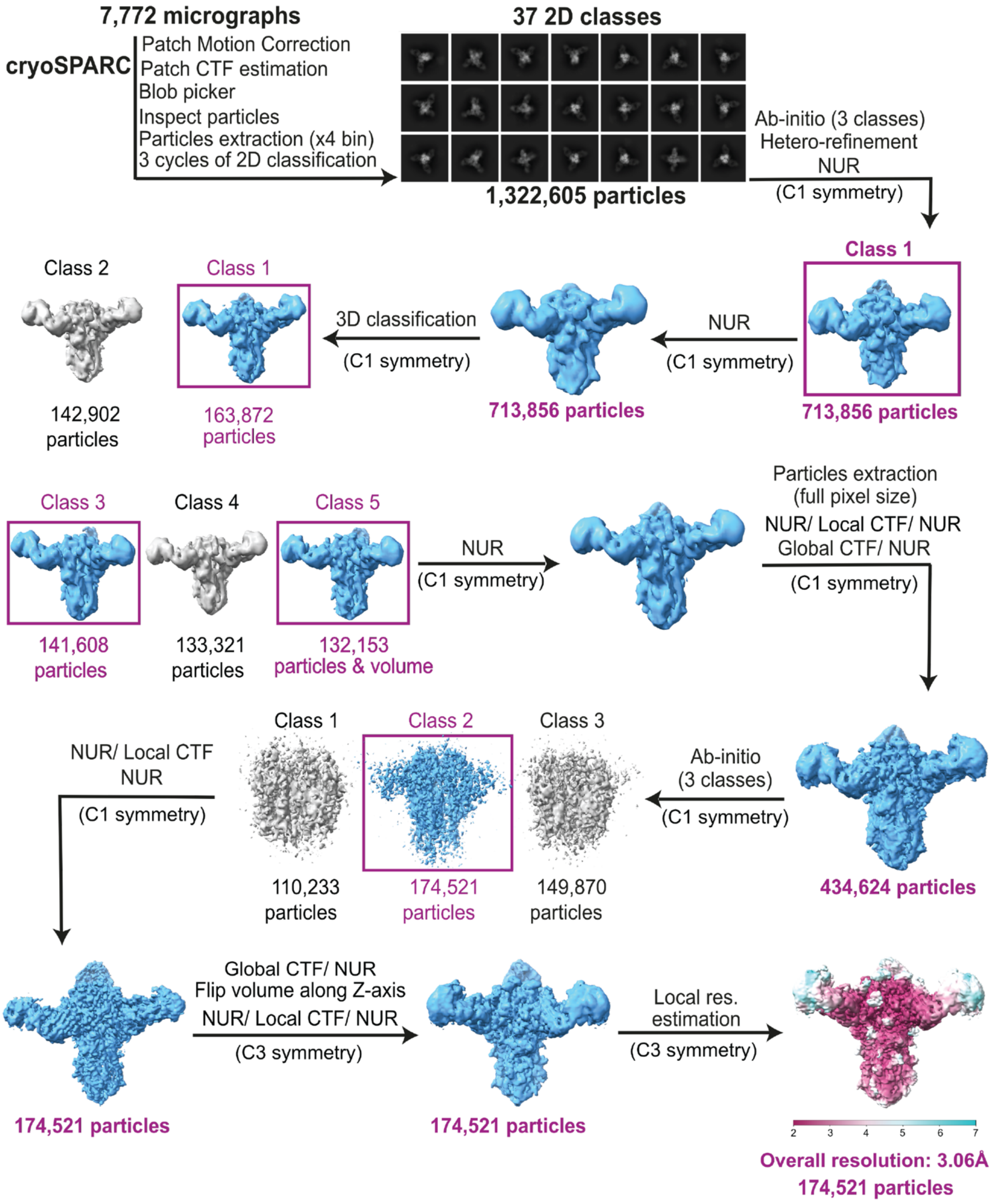


**Supplementary Fig. 5: Cryo-EM processing scheme for ADI-85647 in complex with A/Darwin/6/2021.**

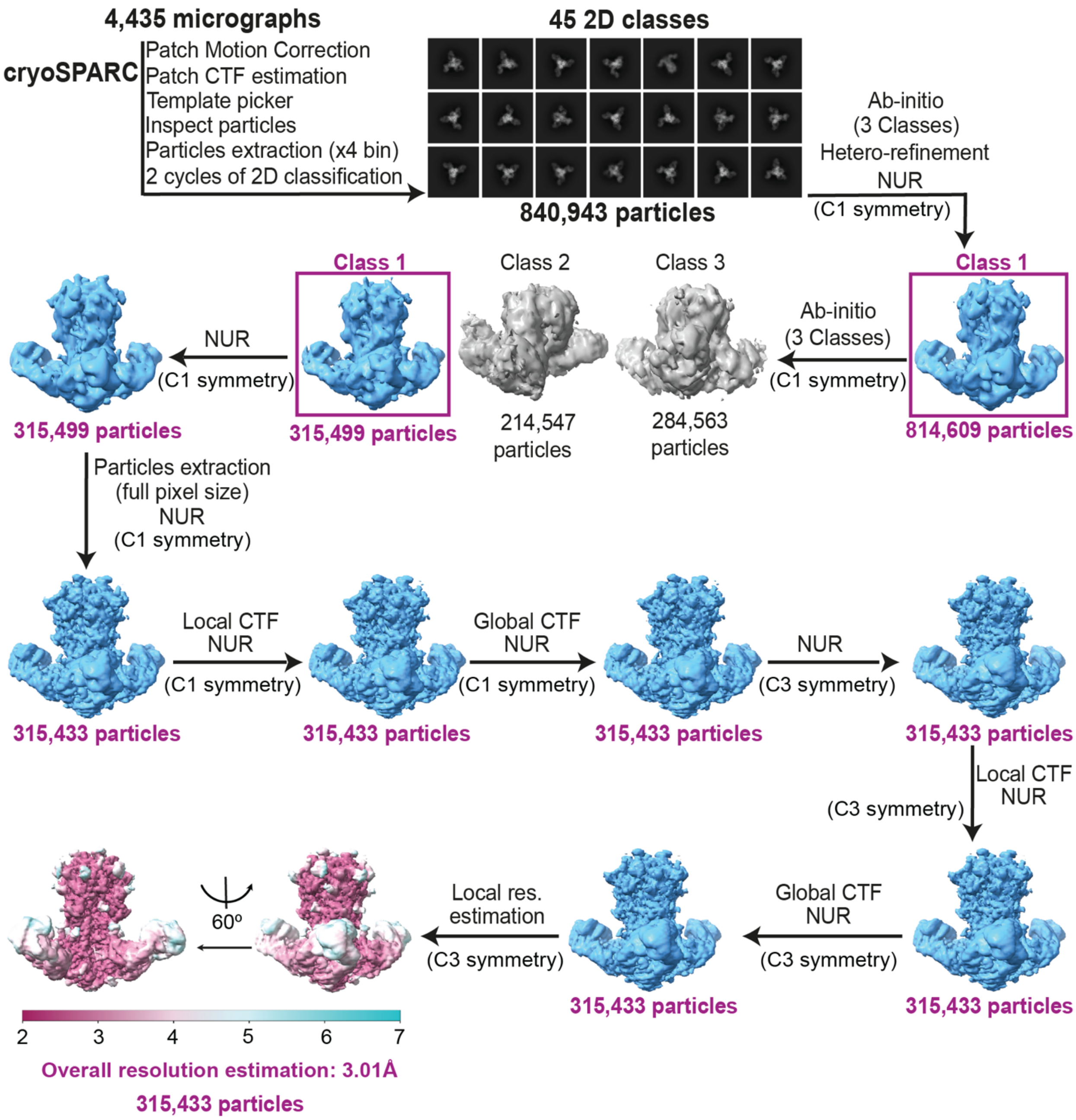


**Supplementary Fig. 6: Cryo-EM processing scheme for ADI-85666 in complex with A/Darwin/6/2021.**
